# Evaluating the efficacy of multi-echo ICA denoising on model-based fMRI

**DOI:** 10.1101/2022.06.29.498113

**Authors:** Adam Steel, Brenda D. Garcia, Edward H. Silson, Caroline E. Robertson

## Abstract

fMRI is an indispensable tool for neuroscience investigation, but this technique is limited by multiple sources of physiological and measurement noise. These noise sources are particularly problematic for analysis techniques that require high signal-to-noise ratio for stable model fitting, such as voxel-wise modeling. Multi-echo data acquisition in combination with echo-time dependent ICA denoising (ME-ICA) represents one promising strategy to mitigate physiological and hardware-related noise sources as well as motion-related artifacts. However, most studies employing ME-ICA to date are resting-state fMRI studies, and therefore we have a limited understanding of the impact of ME-ICA on task or model-based fMRI paradigms. Here, we addressed this knowledge gap by comparing data quality and model fitting performance on data acquired during a visual population receptive field (pRF) mapping paradigm (N=13 participants) after using one of three preprocessing procedures: ME-ICA, optimally combined multi-echo data without ICA-denoising, and typical single echo processing. As expected, multi-echo fMRI improved temporal signal-to-noise compared to single echo fMRI, with ME-ICA amplifying the improvement compared to optimal combination alone. However, unexpectedly, this boost in temporal signal-to-noise did not directly translate to improved model fitting performance: compared to single echo acquisition, model fitting was only improved after ICA-denoising. Specifically, compared to single echo acquisition, ME-ICA resulted in improved variance explained by our pRF model throughout the visual system, including anterior regions of the temporal and parietal lobes where SNR is typically low, while optimal combination without ICA did not. ME-ICA also improved reliability of parameter estimates compared to single echo and optimally combined multi-echo data without ICA-denoising. Collectively, these results suggest that ME-ICA is effective for denoising task-based fMRI data for modeling analyses and maintains the integrity of the original data. Therefore, ME-ICA may be beneficial for complex fMRI experiments, including task fMRI studies, voxel-wise modeling, and naturalistic paradigms.

## Introduction

Functional MRI data is powerful tool for investigating neural activity in the human brain, providing a window into brain organization (Avena-Koenigsberger et al., 2017; Bassett and Bullmore, 2006, 2017; Bassett and Sporns, 2017; Buckner et al., 2011; Bullmore and Sporns, 2012; Busch et al., 2022; Feilong et al., 2021; Gomez et al., 2019a; Gordon et al., 2017; Gratton et al., 2018; Grill-Spector and Weiner, 2014; Huntenburg et al., 2018; Kanwisher et al., 1997; Laumann et al., 2015; Margulies et al., 2016; Murphy et al., 2018; Power et al., 2011; Thomas Yeo et al., 2011) and neural computations (Allen et al., 2021; Baldassano et al., 2017; Breedlove et al., 2020; Caucheteux and King, 2022; Chang et al., 2021; Constantinescu et al., 2016; Doeller et al., 2010; Gomez et al., 2019a; Güçlü and van Gerven, 2015; Hasson et al., 2008; Honey et al., 2012; Huth et al., 2016, 2012; Kay et al., 2015a; Kriegeskorte et al., 2008; Lescroart and Gallant, 2019; Popham et al., 2021; Sha et al., 2015; Wager et al., 2013). However, the contribution of nonneuronal noise, such as motion, heart rate, respiration, and hardware-related artifacts, severely impacts the quality of fMRI data (Bright and Murphy, 2017; Caballero-Gaudes and Reynolds, 2017; Friston et al., 1996; Liu, 2016). As such, optimizing data acquisition and preprocessing/denoising is critically important for ensuring accurate and reproducible results in all fMRI studies.

One promising data acquisition and preprocessing procedure is multi-echo fMRI (Poser et al., 2006; Posse, 2012) combined with echo-time (TE) dependent ICA denoising (hereafter referred to collectively as ME-ICA (Kundu et al., 2017, 2012)). The ME-ICA procedure is described in detail elsewhere (DuPre et al., 2019; Evans et al., 2015; Kundu et al., 2017, 2012), but in brief, ME-ICA involves two steps. First, during acquisition, researchers acquire multiple TEs at each repetition time (TR) (Kundu et al., 2017; Poser et al., 2006). Second, during preprocessing researchers use ICA to decompose the fMRI signal into multiple sources (components). These components are then classified as signal and noise by leveraging the differential decay rate of BOLD-like and non-BOLD signals across TEs (Evans et al., 2015; Kundu et al., 2012).

In principle, ME-ICA helps to resolve two central limitations of single echo fMRI: i) heterogenous signal quality across the echo planar image (EPI) volume due to regional variation in the optimal TE (Kundu et al., 2017; Poser et al., 2006) and ii) noise sources that are not easily differentiable from signal (Kundu et al., 2017). First, because multiple TEs are collected, an optimal TE can be calculated for all voxels and synthesized by a weighted combination of the echoes (Kundu et al., 2012; Poser et al., 2006; Turker et al., 2021). This benefit is most apparent in regions impacted by susceptibility artifacts, like the orbital frontal cortex and lateral temporal lobes (Deichmann et al., 2003; Weiskopf et al., 2007, 2006), where including short echo times greatly improves signal (Kundu et al., 2017; Poser et al., 2006). Second, ME-ICA denoising offers a data-driven method for identifying and removing various noise sources from the neural signal. ME-ICA is particularly effective for removing physiological noise such as cardiac and respiratory signals, which can be challenging to model effectively (DuPre et al., 2021, 2019; Evans et al., 2015; Kundu et al., 2012; Spreng et al., 2019). These benefits make multi-echo a promising strategy for data acquisition, particularly when combined with TE-dependent ICA denoising.

How effective is the ME-ICA procedure at denoising fMRI data? Studies evaluating ME-ICA have focused largely on its application in resting-state fMRI, where the underlying neural signal cannot be explicitly modelled. In these studies, ME-ICA reliably identifies known noise sources, including motion-related artifacts, physiological signals, and thermal noise for removal (Kundu et al., 2012). After ME-ICA denoising, studies typically report significantly greater correlation values among regions within known functional networks (Cohen et al., 2021; Kundu et al., 2012; Lynch et al., 2020; Olafsson et al., 2015) and in difficult to image brain areas like the locus coereleus (Turker et al., 2021). However, because no ‘ground truth’ signal exists in resting-state data, it is challenging to quantify denoising success. Therefore, resting-state is undesirable for evaluating denoising performance for model-based fMRI analysis (e.g., task-fMRI).

Few studies have evaluated the impact of ME-ICA on task-based fMRI. In one study, the authors showed participants grating stimuli with slowly varying stimulus contrast (Evans et al., 2015). They found that ME-ICA enabled detection of this drifting neural signal, which was not possible using single echo processing (Evans et al., 2015). Similarly, a second study found that ME-ICA led to more consistent cardiovascular reactivity mapping during breath-hold challenges (Moia et al., 2021). While these studies demonstrate unique advantages of ME-ICA, they did not report measures of data quality, such as signal-to-noise or model fit, so it is not clear whether this benefit would generalize to other task paradigms or study designs. More recently, a study found that ME-ICA improved signal quality when participants performed a verbal report task, which causes unavoidable head motion. However, these authors did not quantitatively evaluate data quality, but instead evaluated denoising success based upon confirmation of their hypothesis after ME-ICA was applied but not before (Gilmore et al., 2022). Finally, one study evaluated the impact of ME-ICA on effect size estimates during a mentalizing task and found that ME-ICA offered a significant benefit to detect univariate activation (Lombardo et al., 2016). While this result was promising (Lombardo et al., 2016), it is unclear whether this benefit translates to more complex voxel-wise models, as well as how ME-ICA affects reliability of parameter estimates. As such, presently it is unclear whether ME-ICA can recover task-related signals and classify them as BOLD-like in the context of complex model-bsaed fMRI experiments (Kundu et al., 2012), or whether these signals might be erroneously discarded, or, worse, propagated across the brain because of biased retention of only task-like signal components, potentially leading to false-positive activation.

Here, we addressed this knowledge gap by quantifying how ME-ICA affects model-based fMRI analyses by comparing ME-ICA to both a minimally pre-processed single echo pipeline and optimally combined multi-echo fMRI without ICA denoising. We were specifically interested in comparing ME-ICA versus single echo and optimal combination with respect to i) data quality, ii) model fitting performance, and iii) reliability of model parameter estimates. To this end, we leveraged the well-studied population receptive field (pRF) mapping paradigm (Amano et al., 2009; Dumoulin et al., 2018; Dumoulin and Wandell, 2008; Groen et al., 2022; Harvey and Dumoulin, 2011; Kay et al., 2015b, 2013; Klein et al., 2014; Larsson and Heeger, 2006; Lerma-Usabiaga et al., 2020; Samuel Schwarzkopf et al., 2014; Sheremata and Silver, 2015; Silson et al., 2016, 2015; Sprague and Serences, 2013; Takemura et al., 2012; van Dijk et al., 2016; Wandell et al., 2007; Wandell and Winawer, 2015; Winawer et al., 2010). In brief, pRF mapping involves systematic spatiotopic stimulation of the visual field to find the optimal visual receptive field for each voxel in the brain (Figure 1) (Dumoulin and Wandell, 2008). The pRF paradigm is ideal for evaluating the impact of ME-ICA for several reasons (Lerma-Usabiaga et al., 2020). First, the spatial distribution of retinotopic coding in the brain is well-described, so this prior knowledge can serve as a basis for evaluating the impact of ME-ICA. Second, because pRF mapping relies on model fitting, the impact of denoising can be evaluated by comparing variance explained (R^2^) across the protocols. Third, because three pRF parameters (position (X, Y) and size (sigma)) are estimated from the data, the impact of each protocol on reliability can be quantified by comparing the parameter estimates from distinct runs of data. To preview our results, we found that ME-ICA significantly improved data quality, model fit performance, and parameter estimate reliability compared to both single echo data and optimal combination alone.

**Figure 1.**
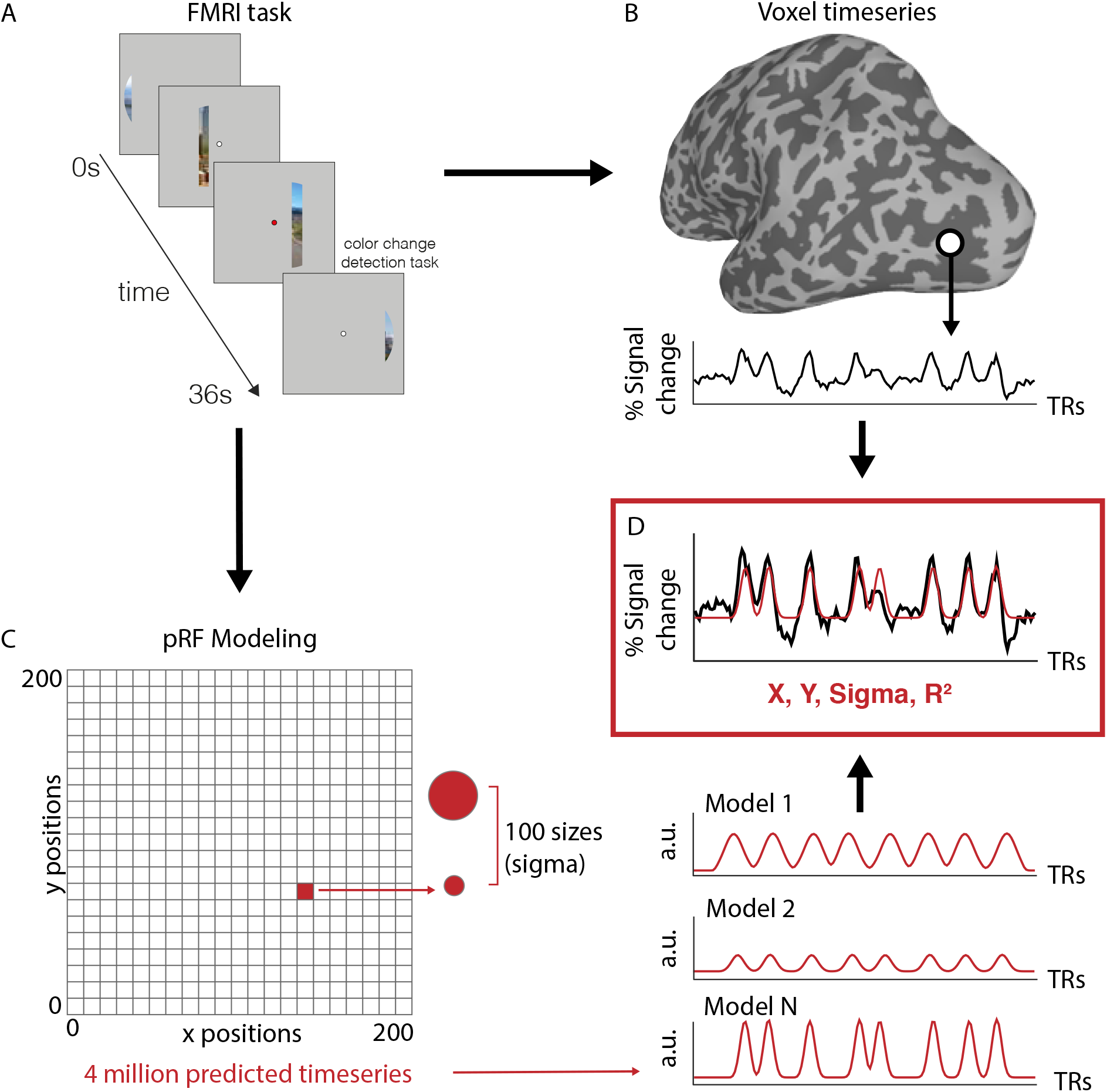
Population receptive field modeling paradigm. A. Task schematic for pRF mapping: Scene images were flashed through a bar aperture that traversed the visual field. A single sweep across the visual field took 36 s and consisted of 18 equal time (2s) and width instances of the aperture. In each run, the aperture completed eight sweeps (2 orientations, 4 directions). Participants were required to maintain fixation and indicate the detection of a color change at fixation via button press. Over an entire sweep, 90 scene images (5 × 18 aperture positions) were presented at random without replacement, guaranteeing that no scene was presented twice within a sweep. This results in a measured timeseries at each fMRI voxel (B). C. To determine the population receptive field for each voxel, a synthetic timeseries is generated for 400 locations in the visual field (200 x and y positions), and 100 sizes (sigma). This results in 4 million possible timeseries that are fit to each voxel’s activity (D). This fitting procedure is done separately for ME-ICA, optimally combined, and single echo data. a.u.: Arbitrary units.

## Methods

### Participants

We recruited 13 participants (10 females, mean age=23.23 ± 3.5 std) for this study. Participants had normal or corrected-to-normal vision, were not colorblind, and were free from neurological or psychiatric conditions. Written consent was obtained from all participants in accordance with the Declaration of Helsinki with a protocol approved by the Dartmouth College Institutional Review Board and Committee for Protection of Human Subjects (CPHS).

### Retinotopy

To map population receptive fields (pRFs), we used a paradigm adapted from Silson et al., 2015 (scene pRF mapping). In brief, we presented portions of scene images through a bar aperture that moved in a stepwise fashion through a circular field (7.3° visual angle). During each 36 s sweep, the bar aperture took 18 evenly spaced steps every 2 s (1 TR). The bar made eight passes in each run (four orientations, two directions: L-R, BR-TL, T-B, BL-TR, R-L, TL-BR, B-T, and TR-BL; L: left, R: right, B: Bottom, T: top). During each bar step (1 TR), we rapidly presented five scene fragments (400 ms per image). All 90 possible scene images were displayed once per sweep, reducing the likelihood that participants mentally “fill in” the underlying image. To ensure fixation, participants performed a color-detection task at fixation, indicating when the fixation dot changed from white to red via button press (semi-random, approximately 2 color changes per sweep). Six runs of pRF data were collected from each participant.

### FMRI

#### MRI Acquisition

All data were collected at Dartmouth College on a Siemens Prisma 3T scanner (Siemens, Erlangen, Germany) equipped with a 32-Channel head coil. Images were transformed from dicom to nifti format using dcm2niix (v1.0.20190902) (Li et al., 2016).

#### T1 image

For registration purposes, a high-resolution T1-weighted magnetization-prepared rapid acquisition gradient echo (MPRAGE) imaging sequence was acquired (TR = 2300 ms, TE = 2.32 ms, inversion time = 933 ms, Flip angle = 8°, FOV = 256 × 256 mm, slices = 255, voxel size = 1 × 1 × 1 mm). T1 images segmented and surfaces were generated using Freesurfer (Dale et al., 1999; Fischl, 2012; Fischl et al., 2002) (version 6.0) and SUMA (Saad and Reynolds, 2012). Anatomical data were aligned to the fMRI data using AFNI’s (Cox, 1996) align_epi_anat.py and @SUMA_AlignToExperiment (Saad and Reynolds, 2012).

#### Functional MRI Acquisition

FMRI data were acquired using a multi-echo T2*-weighted sequence. The sequence parameters were: TR=2000 ms, TEs=[14.6, 32.84, 51.08], GRAPPA factor=2, Flip angle=70°, FOV=240 x 192 mm, Matrix size=90 × 72, slices=52, Multi-band factor=2, voxel size=2.7 mm isotropic. The initial two frames of data acquisition were discarded by the scanner to allow the signal to reach steady state. The full task comprised 144 timepoints.

#### Preprocessing

Multi-echo data processing was implemented based on the multi-echo preprocessing pipeline from afni_proc.py in AFNI (version 21.3.10 Trajan) (Cox, 1996). Signal outliers in the data were attenuated (3dDespike) (Jo et al., 2013). Motion correction was calculated based on the second echo, and these alignment parameters were applied to all runs. For the single echo procedure, we considered only the middle echo. For optimally combined and ME-ICA denoised procedures, the optimal combination of the three echoes was calculated, and the echoes were combined to form a single, optimally weighted timeseries (T2smap.py, distributed with tedana.py (DuPre et al., 2021, 2019)). ME-ICA was then performed for the ME-ICA denoised data (see below).

Following denoising, all images (single echo, optimally combined, and ME-ICA denoised) were blurred with a 5 mm gaussian kernel in the volume (3dBlurInMask), and signals were normalized to percent signal change. No censoring based on motion was applied.

#### Multi-echo ICA

The ME-ICA data were denoised using TE-dependent multi-echo ICA denoising (tedana.py, version 0.0.1 (DuPre et al., 2021, 2019; Evans et al., 2015; Kundu et al., 2012)). In brief, PCA was applied, and thermal noise was removed using the Kundu decision tree method. Subsequently, data was decomposed using ICA, and the resulting components were classified as signal and noise based on the known properties of the BOLD versus noise on the T2* signal decay. Components classified as noise were discarded using *tedana*’s automated classification with accuracy confirmed via visual inspection. The remaining components were recombined to form the denoised timeseries.

#### pRF model

Our goal was to determine the impact of multi-echo fMRI on encoding model fit and parameter reliability. To this end, we performed several analyses that differed in the number of runs averaged together in time to constitute the timeseries fit by the pRF model. Averaging runs in the time domain is typical in pRF mapping studies to increase signal-to-noise of data prior to model fitting (Silson et al., 2016, 2015). The pRF model implementation used for all analyses is described below. Critically, unless otherwise specified, data presented considered all six runs averaged together in time.

Data were analyzed using the pRF implementation in AFNI. The model estimates pRFs using three parameters: center position (X and Y) and a size (sigma). Center positions X and Y are sampled on a cartesian grid with 200 samples across the width and height of the screen, and 100 evenly spaced FWHM varying from 0 to half of the screen width constitute possible pRF sizes (sigma). These result in 4 million possible pRFs for which the timeseries (i.e, bar positions over time) are estimated and convolved with the hemodynamic response function. We then find the best fit timeseries for each voxel by minimizing the least-squares error of the predicted versus actual timeseries (using both Simplex and Powell optimization algorithms). The resulting output contains the best X, Y, and sigma (pRF size) values for each voxel, as well as the explained variance (R^2^). PRF model fitting was conducted in each subject’s original volume space.

FMRI data were mapped to the surface after pRF model fitting using AFNI’s 3dVol2Surf.

### ROI definitions

We considered three ROIs known to have retinotopic response properties (Dumoulin and Wandell, 2008; Silson et al., 2015) to assess model performance in low and high-level visual areas.

For low level areas we considered early visual cortex, which we defined anatomically using Glasser parcellation (Glasser et al., 2016) (visual areas 1-3) defined on the SUMA standard mesh (std.141.) (Argall et al., 2006; Saad and Reynolds, 2012).

To evaluate model fits in high-level visual areas, in each subject we independently defined the scene selective areas on the brain’s lateral (occipital place area; OPA (Dilks et al., 2013)) and ventral (parahippocampal place area; PPA (Epstein and Kanwisher, 1998)) surface. These regions were established using the same criterion we used in our prior work (Steel et al., 2021). Participants passively viewed blocks of scene, face, and object images presented in rapid succession (500 ms stimulus, 500 ms ISI). Blocks were 24 s long, and each run comprised 12 blocks (4 blocks/condition). There was no interval between blocks. Participants performed two runs of the scene perception localizer. Scene and face areas were drawn based on a general linear test comparing the coefficients of the GLM during scene versus face blocks. These contrast maps were then transferred to the SUMA standard mesh (std.141) using @SUMA_Make_Spec_FS and @Suma_AlignToExperiment (Argall et al., 2006; Saad and Reynolds, 2012). A vertex-wise significance of p < 0.001 along with expected anatomical locations was used to define the regions of interest (Julian et al., 2012; Steel et al., 2021; Weiner et al., 2018).

### Data analysis and statistics

For ROI-based analyses, surface nodes were considered if they survived the following criterion in all preprocessing procedures:

- R^2^ value greater than 0.1, which is within the typical range for pRF mapping studies (Gomez et al., 2019a, 2019b; Silson et al., 2016, 2015)
- Center position (X and Y value) between −0.95 and 0.95 relative to minimum and maximum of the cartesian grid
- Sigma less than 0.95 of the maximum modeled width

Because of the non-normal distribution of R^2^ and parameter estimates, for vertex-based analyses we employed non-parametric statistical tests (Kruskal-Wallis and Wilcoxen rank-sum tests). For across participant analyses, we used standard parametric tests (ANOVAs).

## Results

### Multi-echo ICA denoising improves temporal signal-to-noise across the brain

We first sought to determine the impact of ME-ICA on data quality during task fMRI. To this end, we quantified the temporal signal-to-noise (tSNR) of the preprocessed timeseries. We divided the signal average (here, timeseries means) by the standard deviation of the noise (here, the residual series after PRF estimation). ME-ICA clearly improved tSNR compared to standard preprocessed optimal combined or single echo data (Figure 2). The average tSNR for ME-ICA processed data approximately was 1.6× and 1.3× greater than the single echo and optimally combined data, respectively (mean±std tSNR: single echo=167.7±44.8, optimally combined=198±42.7, ME-ICA=266.5±55). This improvement was due exclusively to a decrease in the standard deviation (denominator). The optimally combined signal offered a modest improvement compared to single echo. Consistent with the increased signal from the short TE, the improvement from optimal combination was concentrated in the ventral medial prefrontal cortex and lateral temporal lobes.

**Figure 2.**
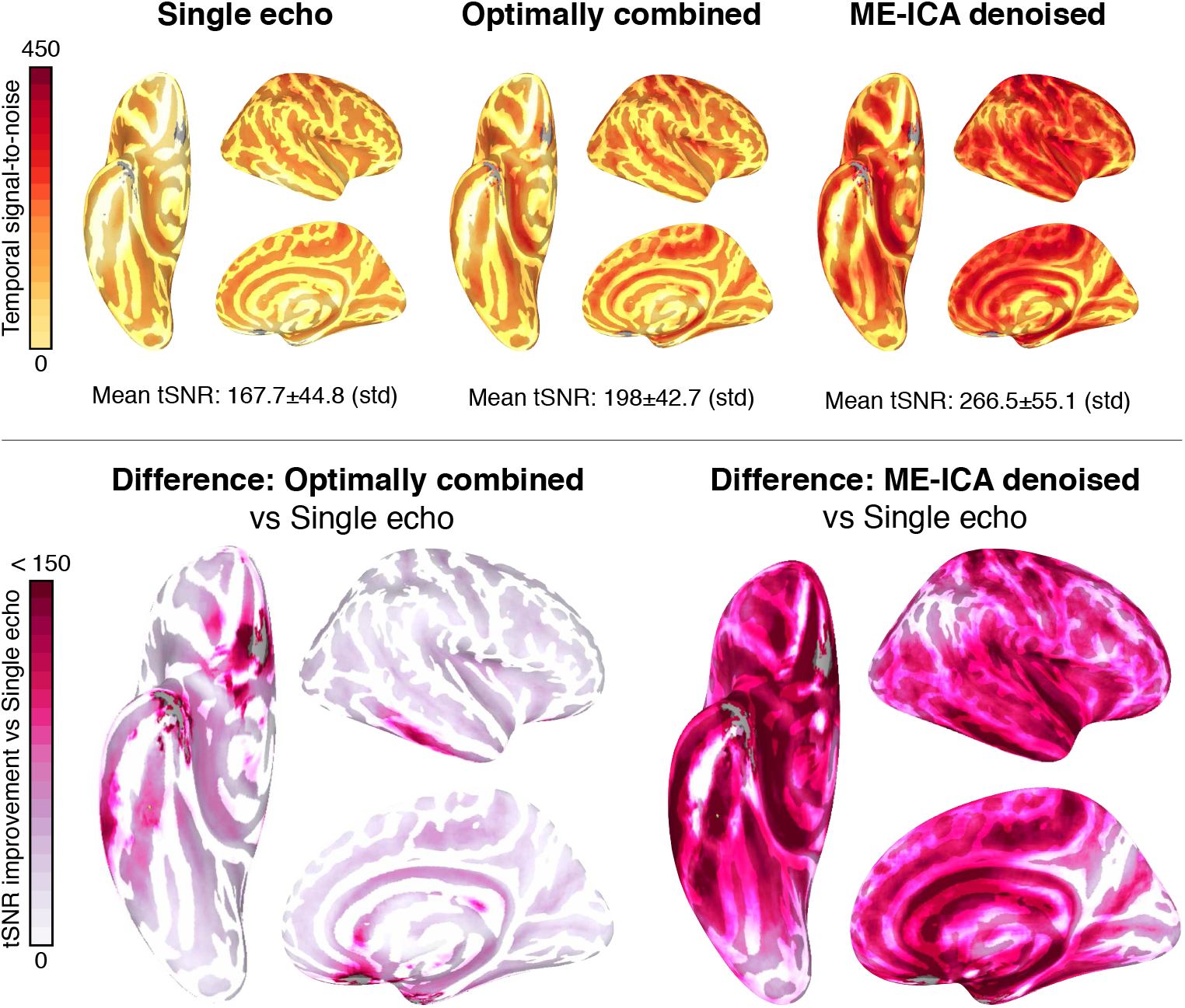
Multi-echo fMRI improves temporal signal-to-noise (tSNR) compared to single echo with minimal preprocessing. We calculated tSNR by taking the mean signal divided by standard deviation of the residuals after pRF model fitting. The ME-ICA procedure significantly improved tSNR compared to single echo and optimal combination alone. Upper panel shows whole brain tSNR values. Lower panel shows difference relatively to single echo for (left) optimal combination and (right) optimal combination + ME-ICA denoising

### Multi-echo ICA denoising improves variance explained by the retinotopic encoding model

The previous result confirmed that ME-ICA denoising and optimal combination with single echo processing improves tSNR compared to single echo during task fMRI. But does the improved tSNR translate to better model fitting? We addressed this question by comparing the R^2^ values between the three processing strategies.

We found a significant improvement in model fitting after ME-ICA compared to single echo and optimally combined data (Figure 3). Interestingly, although optimal combination without ME-ICA denoising yielded improved tSNR, optimal combination alone offered little improvement compared to single echo acquisition.

**Figure 3.**
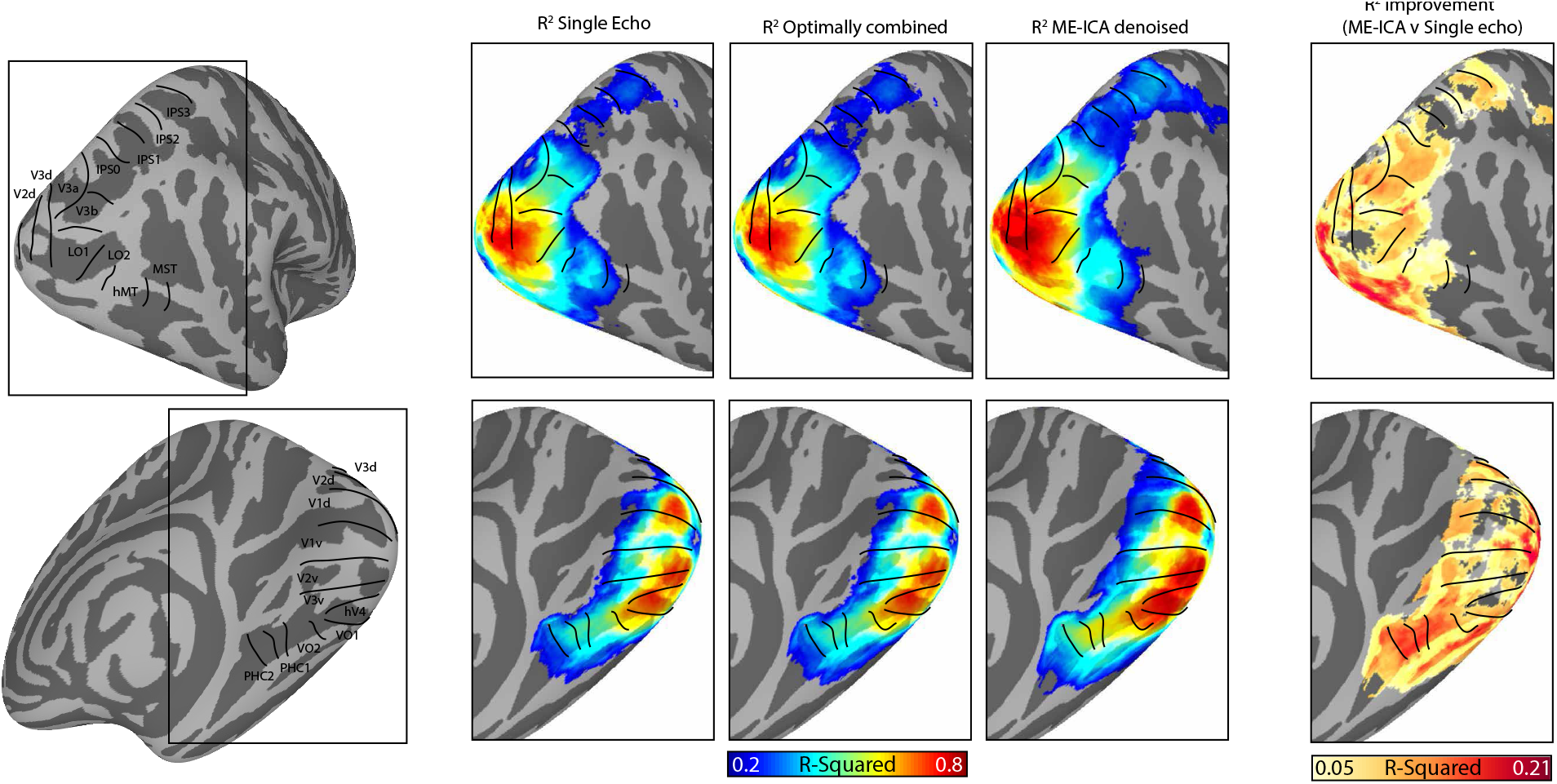
ME-ICA procedure results in significant improvement in pRF model fits across visual cortex. ME-ICA improved R^2^ by as much as 21%, with the largest improvements occuring in ventral temporal cortex. The colorbar is scaled equivalently for R^2^ maps across all preprocessing procedures.

Across the majority of retinotopic cortex, R^2^ was greater for ME-ICA compared to single echo and optimal combination alone, with differences as large as 21%. The most pronounced improvement occurred in ventral occipitotemporal cortex, which we attribute to the tSNR increase afforded by optimal combination and ME-ICA. Importantly, model fits at our R^2^ threshold (0.1) do not extend outside of established retinotopic cortex, suggesting that the model fitting does not arise from an artificial propagation of the task-related signal to non-retinotopic cortex.

This whole-brain analysis offered a coarse, high-level overview of the improvement afforded by ME-ICA denoising. To quantify the improvement more directly, we examined model fits in three visual areas with known retinotopic properties: early visual cortex (V1-V3 defined anatomically based on the Glasser parcellation (Glasser et al., 2016)), as well as two high level visual areas defined functionally in each individual: parahippocampal place area (PPA; (Epstein and Kanwisher, 1998)) on the ventral surface with a upper-field bias, and occipital place area (OPA; (Dilks et al., 2013; Hasson et al., 2003, 2002)) on the lateral surface with a lower-visual field bias (Silson et al., 2015). Example timeseries from voxels in these regions from individual participants are shown in Figure 4, and the distribution of R^2^ values in these areas from all vertices in all individuals is shown in Figure 5. The improvement in R^2^ from ME-ICA is readily apparent in all regions, and the distributions are significantly different (Early visual cortex – left hemisphere: X^2^(2,214607)=5752.29, p<0.0001; right hemisphere: X^2^(2,215610)=5821.19, p<0.0001; OPA – left hemisphere: X^2^(2,8253)=525.62, p<0.0001, right hemisphere: X^2^(2,9422)=527.85, p<0.0001; PPA – left hemisphere: X^2^(2,8070)=737.39, p<0.0001, right hemisphere: X^2^(2,8792)=953.55, p<0.0001). These results demonstrate that ME-ICA denoising improved model fitting success compared to both single echo acquisition and optimal combination without ICA-denoising.

**Figure 4.**
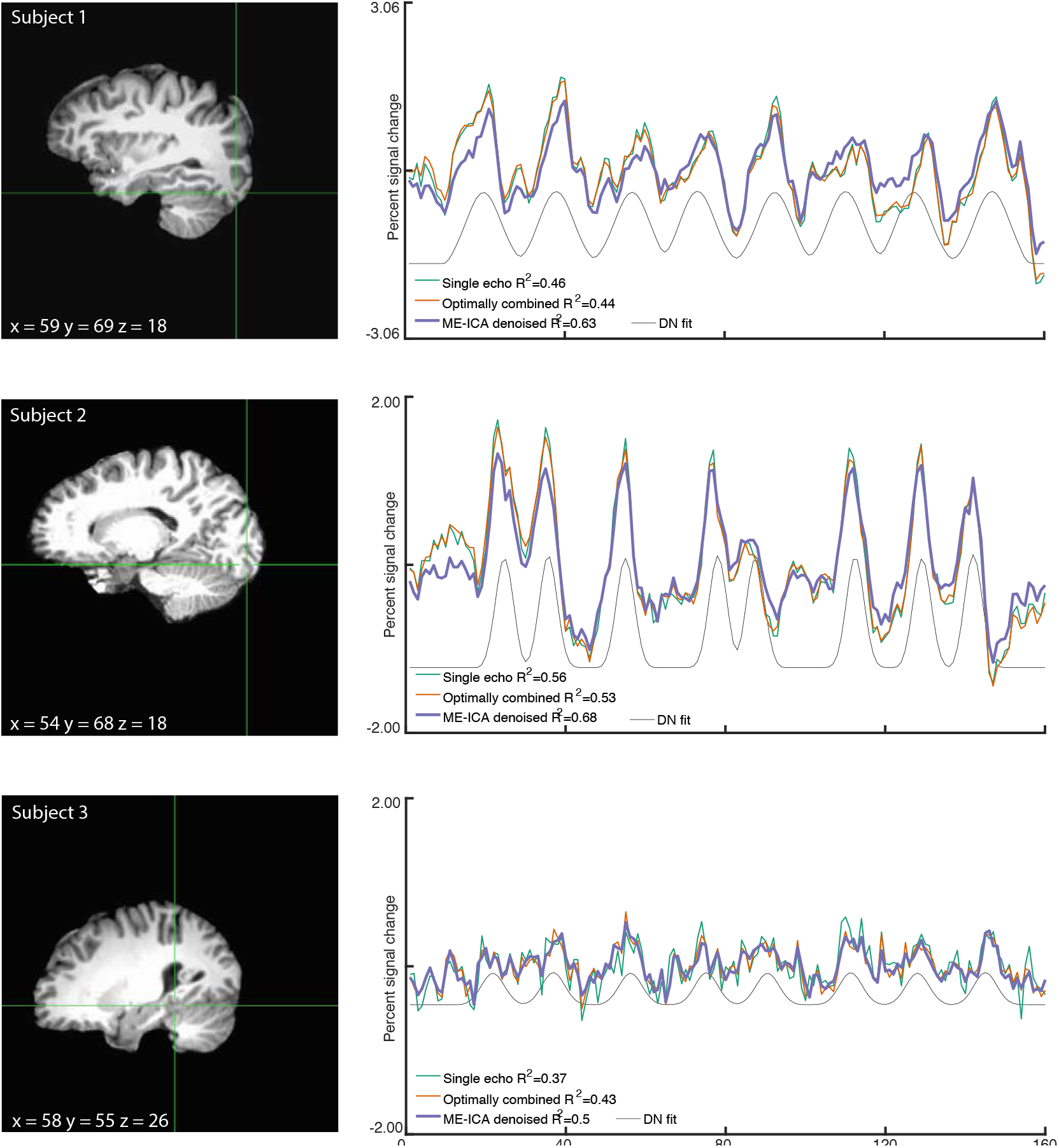
Example voxels timeseries from individual participants. The improvement from ME-ICA appears to result from removed high-frequency noise, which is particularly evident in subject 3 (bottom). Grey line depicts the model fit from the denoised timeseries for reference.

**Figure 5.**
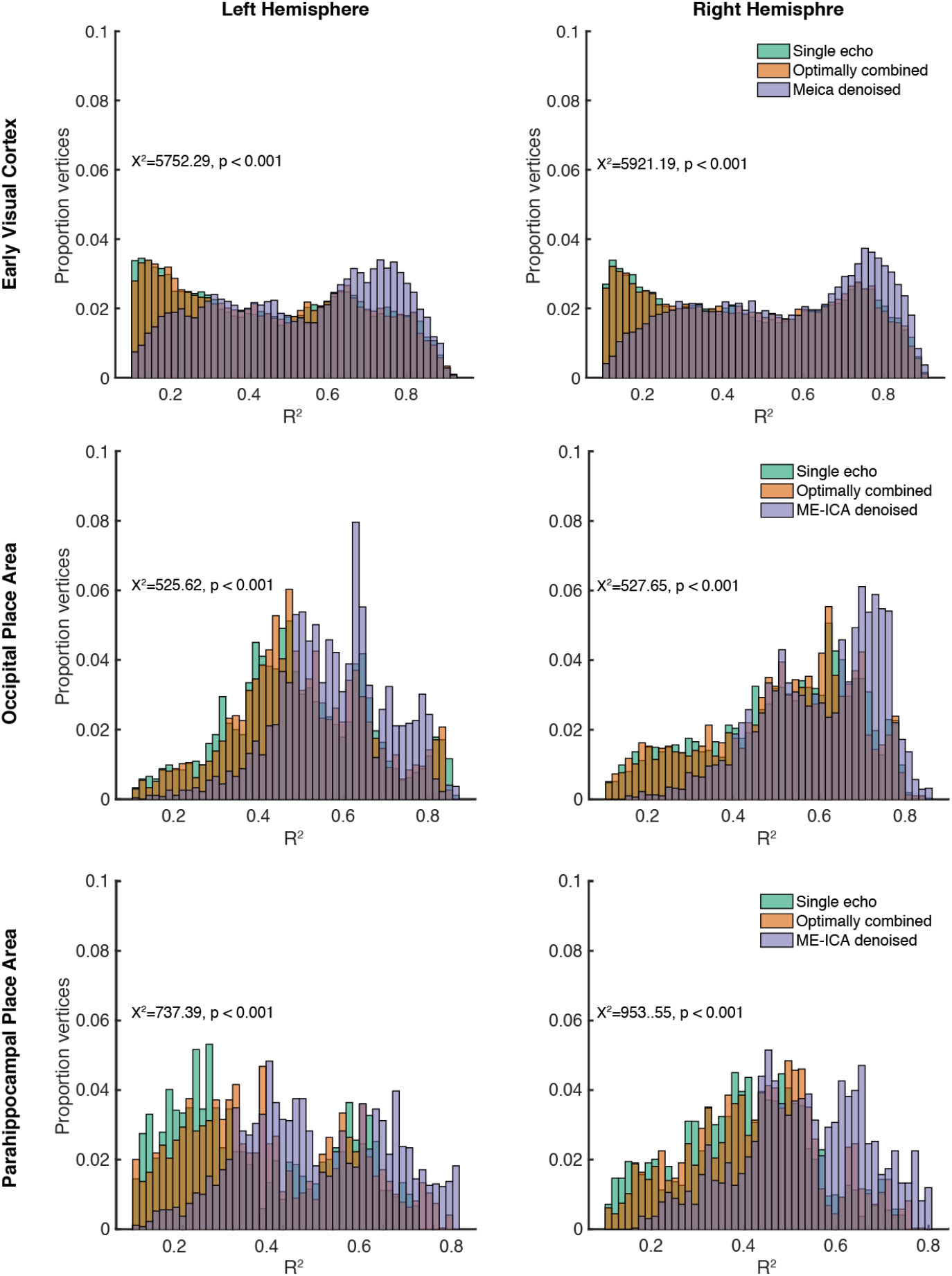
Distribution of R^2^ values from early visual cortex (top), occipital place area (middle), and parahippocampaal place area (bottom) across the data processing procedures. ME-ICA resulted in significantly greater variance explained in all regions.

**Figure 6.**
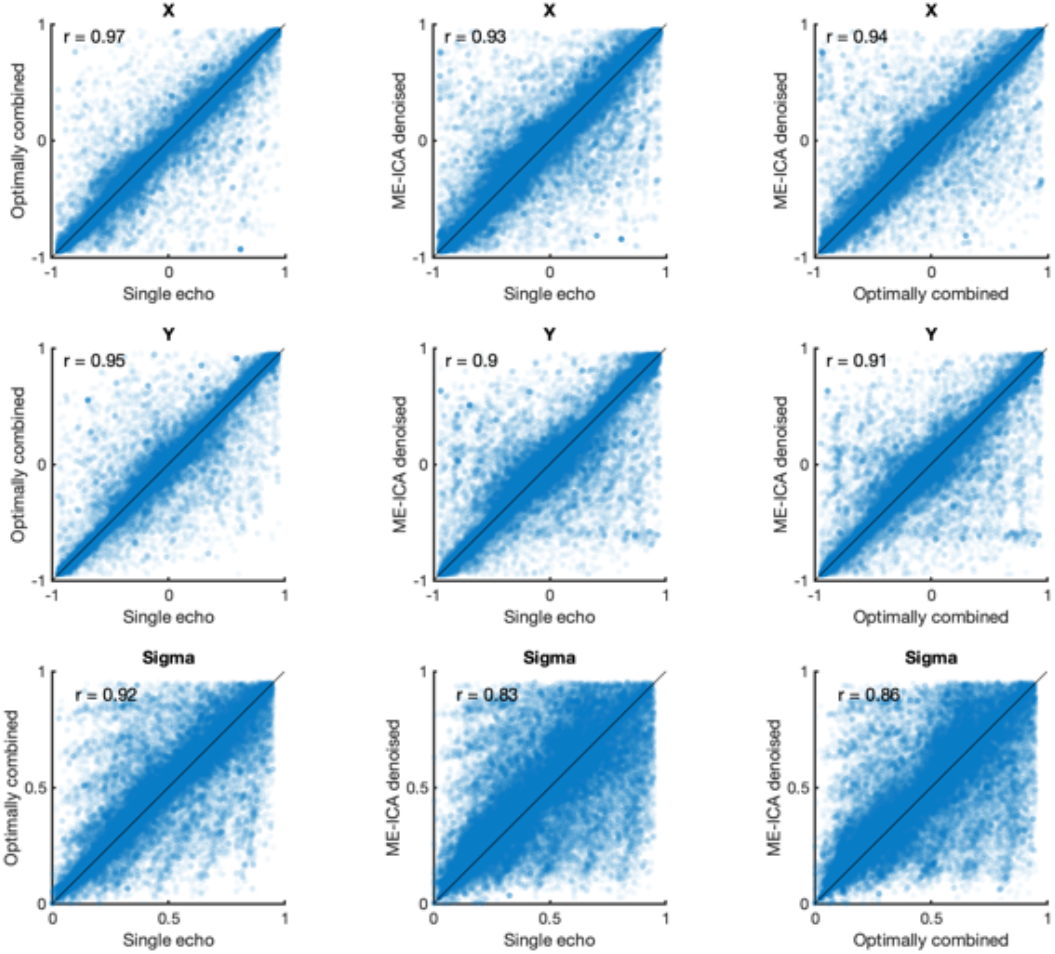
Population receptive field model parameter estimates are highly reliable for early visual cortex. Each plot depicts the vertex-wise correlation between two processing procedures. Data points represent parameter estimates from a single vertex from a single participant. Top row: X parameter, Middle row: Y parameter, Bottom row: Sigma parameter. Black line indicates the unity line.

### ME-ICA increases reliability of parameter estimates with limited data

Our analysis of model fitting suggests that ME-ICA denoising offers substantial improvement in model fitting. However, it is possible that parameter estimates from ME-ICA are not robust if the components do not adequately describe the original signal. We investigated the robustness and reliability in two ways. First, to determine whether ME-ICA denoising preserved the underlying signal, we compared the parameters estimated from ME-ICA denoised data with parameters estimated from the optimally combined and single echo data at every supra-threshold vertex for all subjects. For this analysis, we considered the average of all six data runs. For simplicity, we considered both hemispheres together.

Consistent with ME-ICA preserving the underlying data signal, we found that pRF parameter estimates were highly correlated in all regions of interest across the preprocessing regimes. In all regions, pRF center location estimates were correlated across all preprocessing procedure (Figures 3-5) (X – EVC: SExOC: r(143405) = 0.97; SExDN: r(143405) = 0.93; OCxDN: r(143405) = 0.94; OPA: SExOC: r(5696) = 0.98; SExDN: r(5696) = 0.97; OCxDN: r(5696) = 0.97; PPA: : SExOC: r(5326) = 0.98; SExDN: r(5326) = 0.96; OCxDN: r(5326) = 0.97; Y – EVC: SExOC: r(143405) = 0.95; SExDN: r(143405) = 0.9; OCxDN: r(143405) = 0.91; OPA: SExOC: r(5696) = 0.97; SExDN: r(5696) = 0.96; OCxDN: r(5696) = 0.95; PPA: SExOC: r(5326) = 0.94; SExDN: r(5326) = 0.88; OCxDN: r(5326) = 0.91). Sigma estimates were well-correlated between ME-ICA compared with optimally combined and single echo data for early visual cortex and OPA (Sigma – EVC: SExOC: r(143405) = 0.92; SExDN: r(143405) = .83; OCxDN: r(143405) = 0.86; OPA: r(5696) = .95; SExDN: r(5696) = 0.83; OCxDN: r(5696) = 0.85). PPA showed lower correlation between sigma estimates from ME-ICA with the other procedures (PPA: SExOC: r(5326) = 0.82; SExDN: r(5326) = 0.64; OCxDN: r(5326) = 0.69). This decreased correlation in PPA was due to a shift towards narrower pRF estimates (smaller FWHM) after ME-ICA denoising compared with optimally combined and single echo data, which could suggest greater precision in pRF parameter estimates after ME-ICA denoising (van Dijk et al., 2016).

**Figure 7.**
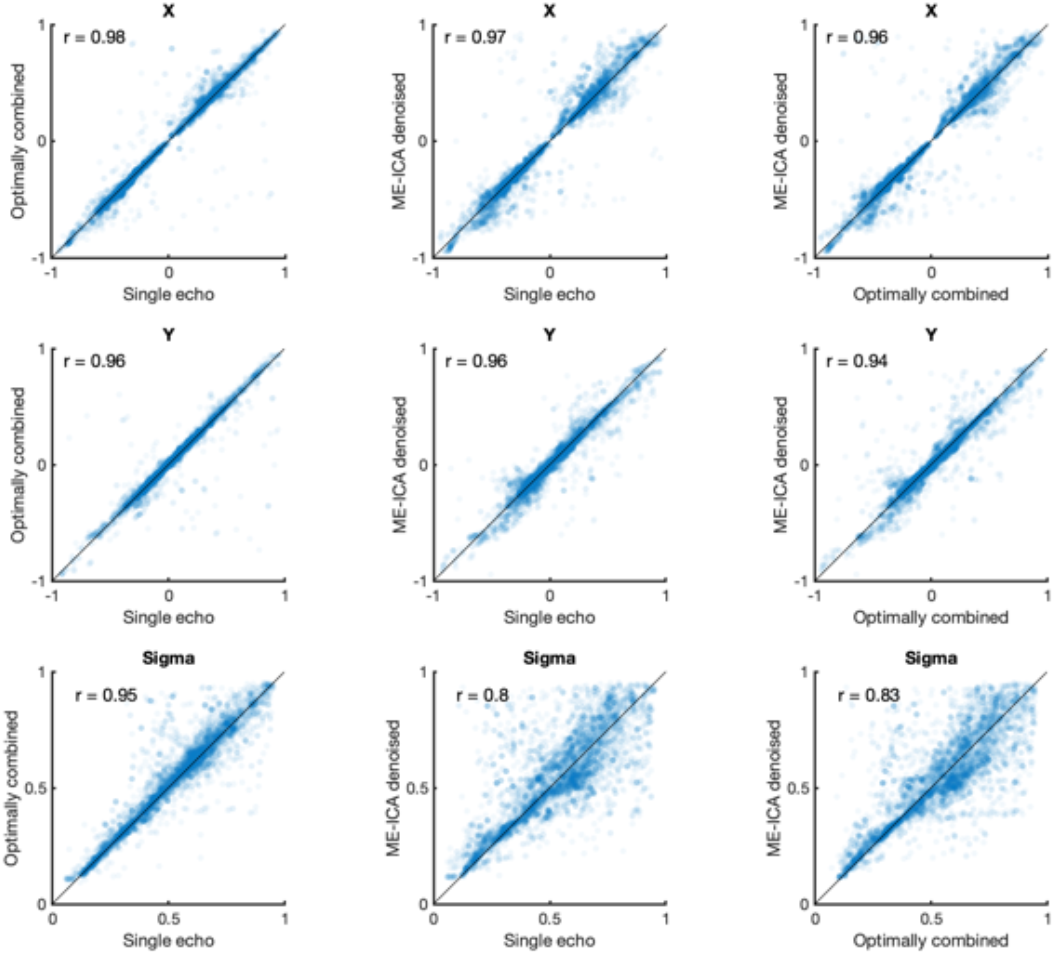
Population receptive field model parameter estimates are highly reliable for OPA. Each plot depicts the vertex-wise correlation between two processing procedures. Data points represent parameter estimates from a single vertex from a single participant. Left column: Single echo × Optimal combination, Middle column, ME-ICA × Single echo, Right column: ME-ICA × Optimal combination. Top row: X parameter, Middle row: Y parameter, Bottom row: Sigma parameter. Black line indicates the unity line.

**Figure 8.**
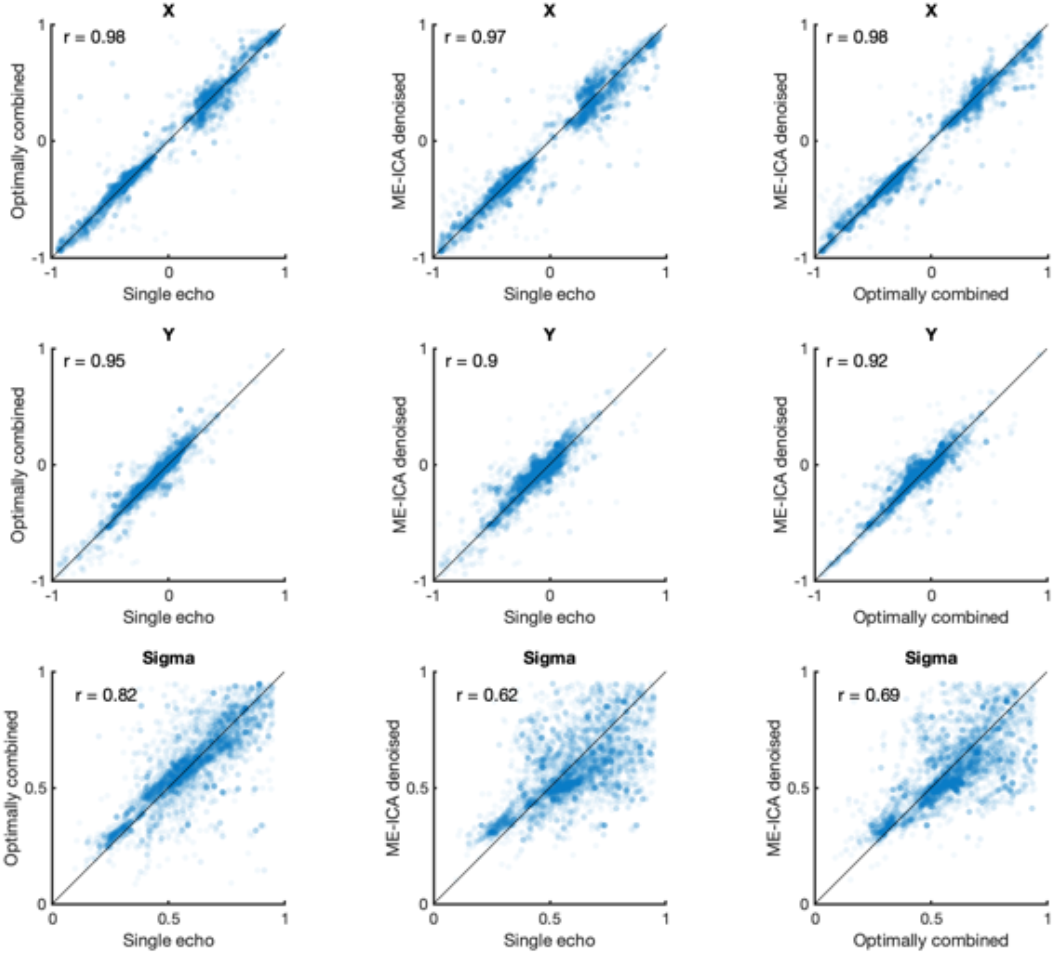
Population receptive field model parameter estimates are highly reliable in PPA. Each plot depicts the vertex-wise correlation between two processing procedures. Data points represent parameter estimates from a single vertex from a single participant. Note that reliability is lower for sigma between ME-ICA and optimally combined/single echo denoising. On average, sigma estimates are lower for ME-ICA, suggesting higher precision sigma estimates. Top row: X parameter, Middle row: Y parameter, Bottom row: Sigma parameter. Black line indicates the unity line.

As a second approach to test parameter estimate reliability, we compared the parameter estimates from each single run of data within a given subject. For each subject, we correlated the run × run parameter estimates for X, Y, and sigma across all vertices in early visual cortex, and compared correlation values across the preprocessing techniques. Higher correlation values indicate greater reliability of parameter estimates. We focused on early visual cortex because each run was estimated independently, so we were unable to meet our R^2^ threshold (0.10) for vertices in PPA and OPA after single echo processing in all subjects. For all three parameters (X, Y, and Sigma) ME-ICA denoising resulted in improved reliability (X – Kruskal-Wallis X^2^(2,1167) = 20.58, p < 0.001; rank sum: OC v SE: z=−0.2, p = 0.83, DN v SE: z = 3.865, p = 0.0001, DN v OC: z = 3.985, p < 0.0001; Y – X^2^(2,1167) = 39.41, p < 0.001; rank sum: OC v SE: z=0.10, p = 0.91, DN v SE: z = 5.5448, p < 0.0001, DN v OC: z = 5.3236, p < 0.0001; Sigma – Kruskal-Wallis X^2^(2,1167) = 27.79, p < 0.001; rank sum: OC v SE: z=0.39, p = 0.695, DN v SE: z = 4.815, p < 0.0001, DN v OC: z = 4.2814, p < 0.0001).

**Figure 9.**
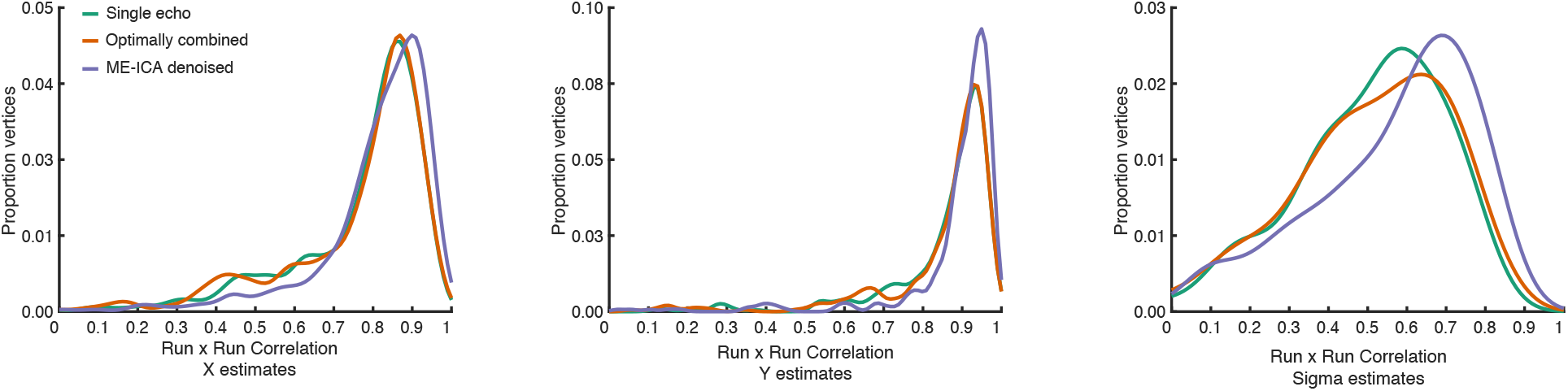
ME-ICA improves reliability of parameter estimates. X, Y, and Sigma estimates were calculated from single runs of pRF mapping. We then correlated these vertex-wise parameter estimates for all vertices within early visual cortex (V1-V3) across all pairs of runs. Reliability of parameter estimates were generally very high. However, ME-ICA resulted in greater reliability than either single echo or optimal combination alone, particularly for sigma.

The above analysis suggested that ME-ICA results in significantly greater reliability with limited data. For our final analysis, we investigated how much R^2^ increased as more data is added. For this analysis, in each subject we averaged together increasing numbers of pRF data runs before pRF model fitting. We then calculated the average R^2^ by the pRF model in early visual cortex at each number of averages. We compared these R^2^ values using a repeated measures ANOVA with number of averages (1-6) and preprocessing procedure (single echo, optimally combined, ME-ICA) as factors. Regardless of preprocessing, R^2^ improved with increasing numbers of runs (Main effect of averages: F(5,334) = 89, p < 0.0001). However, the pRF model explained significantly more variance after ME-ICA compared to the other preprocessing procedures regardless of number of averages (ANOVA: Main effect of preprocessing: F(2,334) = 76.26, p < 0.0001). We found that even with just a single run, ME-ICA resulted in R^2^ roughly equivalent to three runs of optimally combined or single echo data, and just two runs of ME-ICA achieved the same performance as six runs of the other procedures. This finding is consistent with findings investigating precision mapping of functional networks using resting state fMRI (Lynch et al., 2020)).

**Figure 10.**
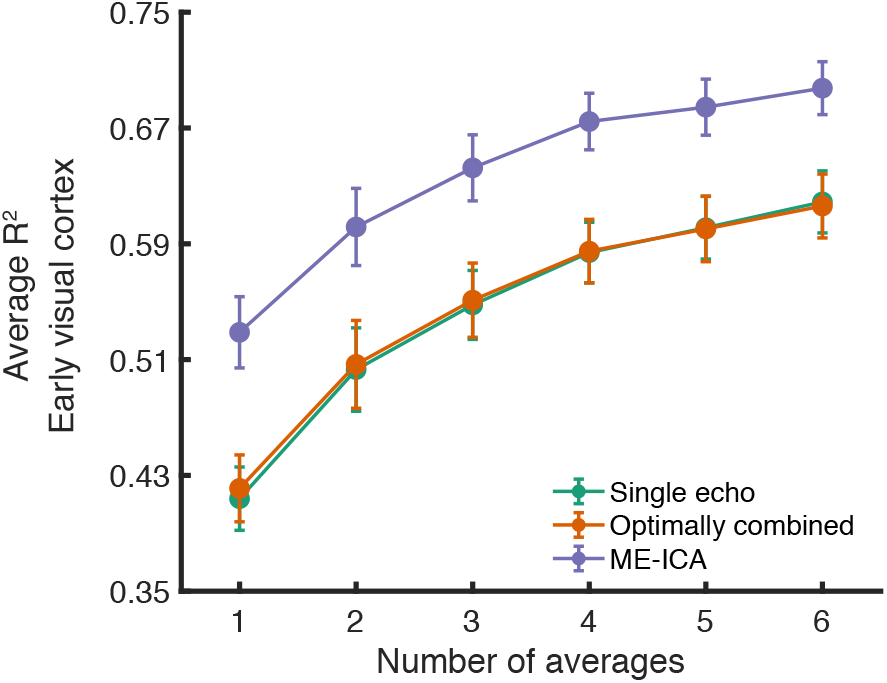
ME-ICA improves model fitting with limited data. In each subject we averaged increasing numbers of runs and calculated the average R^2^ in early visual cortex at each number of averages. ME-ICA benefitted model fitting at all levels of data, with just two runs of data needed to achieve the same R^2^ as six single echo and optimally combined runs.

## Discussion

Here, we investigated the impact of multi-echo acquisition and TE-dependent ICA denoising on model-based fMRI analysis. We found that ME-ICA significantly improved tSNR compared to traditional preprocessing of optimally combined or single echo data resulting from a standard population receptive field (pRF) mapping paradigm (though optimally combining data improved tSNR compared to single echo data, as well). Compared to single-echo or optimal combination without denoising, ME-ICA increased the variance explained by our pRF model throughout the visual system and improved detection of pRFs in difficult to image regions like lateral ventral temporal cortex. ME-ICA also improved reliability of model parameter estimation. Together, these results demonstrate that ME-ICA preprocessing offers a significant benefit for model-based fMRI analyses.

### ME-ICA improves tSNR and model fitting

As expected, we found that multi-echo fMRI acquisition improved tSNR. Both ME-ICA denoised and optimally combined data with typical preprocessing had significantly better tSNR compared to single echo data. The observed improvement was most pronounced in ventral temporal and orbitofrontal cortex because the dropout artifact was mitigated by including the short echo time at acquisition. This finding largely agrees with previous studies that compared results using resting state data (Kundu et al., 2012; Lynch et al., 2020; Turker et al., 2021).

Surprisingly, the tSNR boost did not directly translate to improved model fits. When we compared R^2^ by our retinotopy model across the preprocessing procedures, we found that the model explained significantly more variance in ME-ICA denoised data compared to traditional preprocessing with optimally combined or single echo data. This agrees with a resting state investigation (Boyacioğlu et al., 2015), wherein the authors reported a failure to recover known resting state networks from optimally combined data without ICA denoising (in this case, FIX-ICA (Griffanti et al., 2014; Salimi-Khorshidi et al., 2014)). This finding is important, because one might opt to do multi-echo acquisition with the goal of improving tSNR but not use ME-ICA denoising – for example, this procedure is implemented in one widely used preprocessing framework FMRIPREP (Esteban et al., 2017) (N.B., TE-dependent ICA denoising can be implemented separately on data processed using the FMRIPREP pipeline (DuPre et al., 2021, 2019)). However, our data suggest that model fitting greatly benefits from ME-ICA denoising, and that optimal combination without ME-ICA may only be advantageous compared to single echo acquisition in limited circumstances.

The reason for the dissociation between tSNR and model fitting performance after optimal combination alone is not clear. One possibility is that global noise sources common across all echoes exist in the data, and so acquiring multiple echoes does not equate to the same benefit as independent averages (Power et al., 2017). Given that the average improvement in tSNR did not increase as much as expected (i.e., improvement was less than the square root of number of averages), this seems likely. Additionally, contrast-to-noise ratio is lower in the short echo times. So, despite improved tSNR in areas with shorter optimal TEs, these regions may still have suboptimal contrast-to-noise required for model fitting. Future work should continue examining the relationship between model fitting performance and tSNR.

### ME-ICA improves reliability of model parameter estimation

Beyond R^2^, it is important that preprocessing and analysis choices lead to reproducible outcomes (Botvinik-Nezer et al., 2020; Bowring et al., 2019; Lerma-Usabiaga et al., 2020; Soltysik, 2020). In our study, we leveraged the relative stability of voxel-wise retinotopic tuning to test whether ME-ICA led to similar parameter estimates from independent runs of data (van Dijk et al., 2016). We found that ME-ICA gave highly robust parameter estimates. First, parameters estimated from ME-ICA data were highly correlated to parameters estimated from single-echo data. This result suggests that the ICA procedure does not introduce bias or distort the timeseries signal.

Second, ME-ICA improved the reliability of retinotopy parameter estimation from single runs of fMRI data compared to single echo or optimal combination alone. In addition, we found that ME-ICA preprocessed data led to significantly lower estimates of pRF size compared to traditional preprocessing, suggesting that ME-ICA may increase the precision of pRF estimation (van Dijk et al., 2016).

### Limitations

In the present study, we investigated the impact of ME-ICA denoising compared to a traditional, minimal preprocessing pipeline. Specifically, we sought to confirm that ME-ICA offered improvement in tSNR and model fitting, while not affecting parameter estimates. Our minimal preprocessing included only despiking, motion correction, smoothing, and scaling. This pipeline is similar the pipeline used in our other retinotopy studies (Silson et al., 2016, 2015). This nevertheless represents just one possible choice among many (Caballero-Gaudes and Reynolds, 2017; Moia et al., 2021; Power et al., 2017). Other data-driven denoising techniques using ICA (e.g., hand classification (Griffanti et al., 2017) or automated pipelines such as AROMA-ICA (Pruim et al., 2015) or ICA-FIX (Salimi-Khorshidi et al., 2014)) are highly effective at denoising single echo fMRI data. These techniques can be run on single echo or optimally combined data and thus may be viable preprocessing options if multi-echo acquisition is not possible. However, it is important to note that ICA-based denoising is not a panacea, and some noise sources remain (and can be worsened) after ICA-based denoising strategies, including ME-ICA (Power et al., 2018). Anecdotally, because ICA is non-deterministic, it can fail to converge or fail to produce high-quality components, so it is important to check the output from this technique thoroughly.

Outside of ICA-based techniques, many other denoising strategies can be used (Caballero-Gaudes and Reynolds, 2017). It is common in other fields, like resting state fMRI, to remove high-motion time points from data and to project ‘nuisance regressors’ out of the data prior to analysis (Ciric et al., 2017; Power et al., 2017; Satterthwaite et al., 2013). The effects of these techniques, including CompCorr (Behzadi et al., 2007), ANATICORR (Jo et al., 2010), MotSim (Patriat et al., 2017), and multiple derivatives of motion (Satterthwaite et al., 2013), have been compared in detail elsewhere (Ciric et al., 2017; Moia et al., 2021; Power et al., 2017), and an exhaustive comparison of preprocessing choices is out of the scope of the present work (for a comprehensive review see, (Caballero-Gaudes and Reynolds, 2017)).

Finally, because multi-echo acquisition requires taking multiple EPI volumes in a single TR, multi-echo generally requires a longer TR than optimized single echo acquisition regimes. Relatedly, the sampling duration necessary for super high resolution fMRI imaging can preclude the use of multi-echo fMRI. While these limitations can be overcome using mutli-slice acquisition, in-plane acceleration, or partial Fourier acquisition, these techniques can result in decreased signal-to-noise of the resulting data (Boyacioğlu et al., 2015; Cohen et al., 2021; Tsao and Kozerke, 2012). In the end, the optimal acquisition and preprocessing procedure for any given study depends on the research question, as well as the technical, computational, and hardware resources available.

## Conclusion

To summarize, we found that ME-ICA improves tSNR and model fitting in task-based fMRI data (pRF mapping) both in terms of variance explained and regions implicated. Additionally, our findings suggest that model parameters can be estimated reliably from just a single run of data after ME-ICA processing. Therefore, ME-ICA may be an attractive option for naturalistic fMRI experiments where collecting single runs of fMRI data is common.

## Acknowledgements

We would like to thank Y.B. Choi, Anna Mynick, and Terry Sackett for assistance with data collection and Eneko Uruñuela for helpful discussion. This work was supported by a grant from NVIDIA to CER. AS is supported by the Neukom Institute for Computational Science.

## Notes

Conflict of interest statement: The authors declare no conflict of interest.

### Competing Interest Statement

The authors have declared no competing interest.

### Summary of Updates

Add references

